# The extrinsic incubation period for Zika virus: a Bayesian time delay modelling study

**DOI:** 10.64898/2026.08.05.742962

**Authors:** Anna-Maria Hartner, Lothar H. Wieler, Christopher Irrgang

**Affiliations:** Centre for Artificial Intelligence in Public Health Research, Robert Koch Institute, Berlin, Germany; MRC Centre for Global Infectious Disease Analysis, Jameel Institute, School of Public Health, Imperial College London, London, United Kingdom; Hasso-Plattner-Institute, Digital Health Cluster, University of Potsdam, Potsdam, Germany; Hasso-Plattner-Institute for Digital Health at Mount Sinai, ICAHN School of Medicine, Mount Sinai Hospital, New York, USA

## Abstract

The extrinsic incubation period (EIP), defined as the time between a mosquito acquiring a virus and becoming capable of transmitting it, is a key component of arbovirus transmission and varies with temperature. For Zika virus (ZIKV), empirical estimates of EIP are derived from heterogeneous laboratory studies and are typically analysed without accounting for interval censoring inherent in vector competence experiments. We applied Bayesian interval-censored survival models to pooled individual-level observations from eight published studies, comparing 20 candidate models representing four parametric survival distributions and five temperature-response functions in *Ae. aegypti* and *Ae. albopictus*. Model performance was evaluated using approximate leave-one-out cross-validation. A lognormal survival model with a quadratic temperature response and study-level random intercept provided the best predictive performance. Median EIP declined non-linearly with increasing temperature, from 59.5 days (95% CrI 27.0–124.0) at 20°C to 8.4 days (3.8–17.0) at 32°C in *Ae. aegypti*. Across the temperature range examined, estimated EIPs for *Ae. albopictus* were approximately 1.5-fold longer than those for *Ae. aegypti*. Credible intervals widened at temperature extremes, reflecting between-study heterogeneity and limited data availability. These results provide a statistical framework for estimating temperature-dependent ZIKV EIP while accounting for censoring and uncertainty, supporting improved parameterisation of mechanistic models of arbovirus transmission.

**Author summary:** When a mosquito feeds on an infected host, the virus must replicate and spread to the mosquito’s salivary glands before it can be transmitted. This delay, known as the extrinsic incubation period (EIP), is a key determinant of mosquito-borne disease transmission and varies with temperature. For Zika virus (ZIKV), estimates of EIP come from laboratory studies that differ in experimental design, mosquito species, and viral dose, making them difficult to combine. We analysed data from eight studies using a Bayesian statistical approach that accounts for uncertainty in the time mosquitoes become infectious, because mosquitoes are typically tested only at discrete time points after infection. Our analysis showed that ZIKV EIP decreases non-linearly with increasing temperature, ranging from approximately 60 days at 20°C to approximately 8 days at 32°C in *Aedes aegypti*. Across temperatures, *Aedes albopictus* had consistently longer EIPs than *Aedes aegypti*. Compared with Bayesian EIP models developed for other arboviruses, including dengue, yellow fever, and West Nile virus, ZIKV showed a stronger non-linear temperature response. These estimates and their uncertainty provide improved parameters for models predicting when and where Zika transmission is most likely.

## Introduction

Zika virus (ZIKV) is a mosquito-borne flavivirus transmitted primarily by *Aedes aegypti* and *Aedes albopictus* [1]. The 2015–2016 outbreaks in the Americas demonstrated the potential for rapid ZIKV emergence and highlighted the severe consequences of infection during pregnancy, including congenital Zika syndrome and associated neurological complications [1–3]. Transmission by these vectors is governed by temperature-sensitive biological processes, including mosquito survival, viral replication, and dissemination within the vector [4, 5]. Characterising the temperature dependence of these processes is therefore necessary for improving mechanistic models of ZIKV transmission.

A key component of arbovirus transmission is the extrinsic incubation period (EIP), defined as the time between a mosquito acquiring infection through an infectious blood meal and becoming capable of transmitting the virus [5, 6]. EIP influences the basic reproduction number (*R*_0_) because mosquitoes must survive long enough for the virus to disseminate to the saliva before transmission can occur [7–9]. Temperature is a major determinant of this process, with higher temperatures generally shortening the EIP by accelerating viral replication, although the response varies among mosquito populations, viral strains, and experimental conditions [10–15].

Experimental estimates of EIP are typically obtained from vector competence studies in which mosquitoes are sampled at discrete time points after infection to determine when infectious virus is first detected in saliva [16]. These data present several statistical challenges. Mosquitoes that die before becoming infectious, or that remain non-infectious throughout the observation period, generate right-censored observations, while discrete sampling schedules mean that the exact time of infectiousness is often only partially observed [6, 16]. Despite these characteristics, EIP is frequently summarised using fixed endpoints or point estimates, which do not fully account for censoring or the stochastic nature of the transmission process [6, 16, 17].

Time-to-event methods provide a natural framework for analysing EIP data because they explicitly model the distribution of times to infectiousness while accommodating censored observations [6]. Bayesian survival models further allow information to be shared across heterogeneous experiments, incorporate prior biological knowledge, and propagate uncertainty into downstream estimates. These features are particularly relevant when parameterising mechanistic models of arbovirus transmission, where uncertainty in EIP estimates can influence predictions of transmission under different environmental conditions.

Here, we reanalyse a published ZIKV EIP dataset compiled by Hartner et al. using Bayesian parametric survival models [17]. We compare alternative time-to-event distributions to estimate the temperature dependence of EIP in Aedes aegypti and Aedes albopictus while explicitly accounting for censored observations, providing a statistical framework for incorporating EIP uncertainty into mechanistic models of arbovirus transmission.

## Results

### Data

The final dataset comprised 4,617 censored EIP observations from eight laboratory studies of ZIKV in *Ae. aegypti* and *Ae. albopictus*. Most observations were from *Ae. aegypti* (n = 3,389; 73.4%), and the majority were right-censored (n = 3,545; 76.8%). Experimental temperatures ranged from 16–38°C for *Ae. aegypti* and 18–35°C for *Ae. albopictus*.

### Model Selection

Among the 20 candidate models, the quadratic temperature structure with a lognormal survival distribution (f3 lognormal) achieved the highest predictive performance according to PSIS-LOO (Fig. 1). Model comparisons indicated that temperature-response structure was the dominant determinant of predictive performance: replacing linear temperature terms (f1/f2) with a quadratic temperature response (f3) resulted in a substantial improvement in expected log predictive density (ELPD).

**Fig 1.**
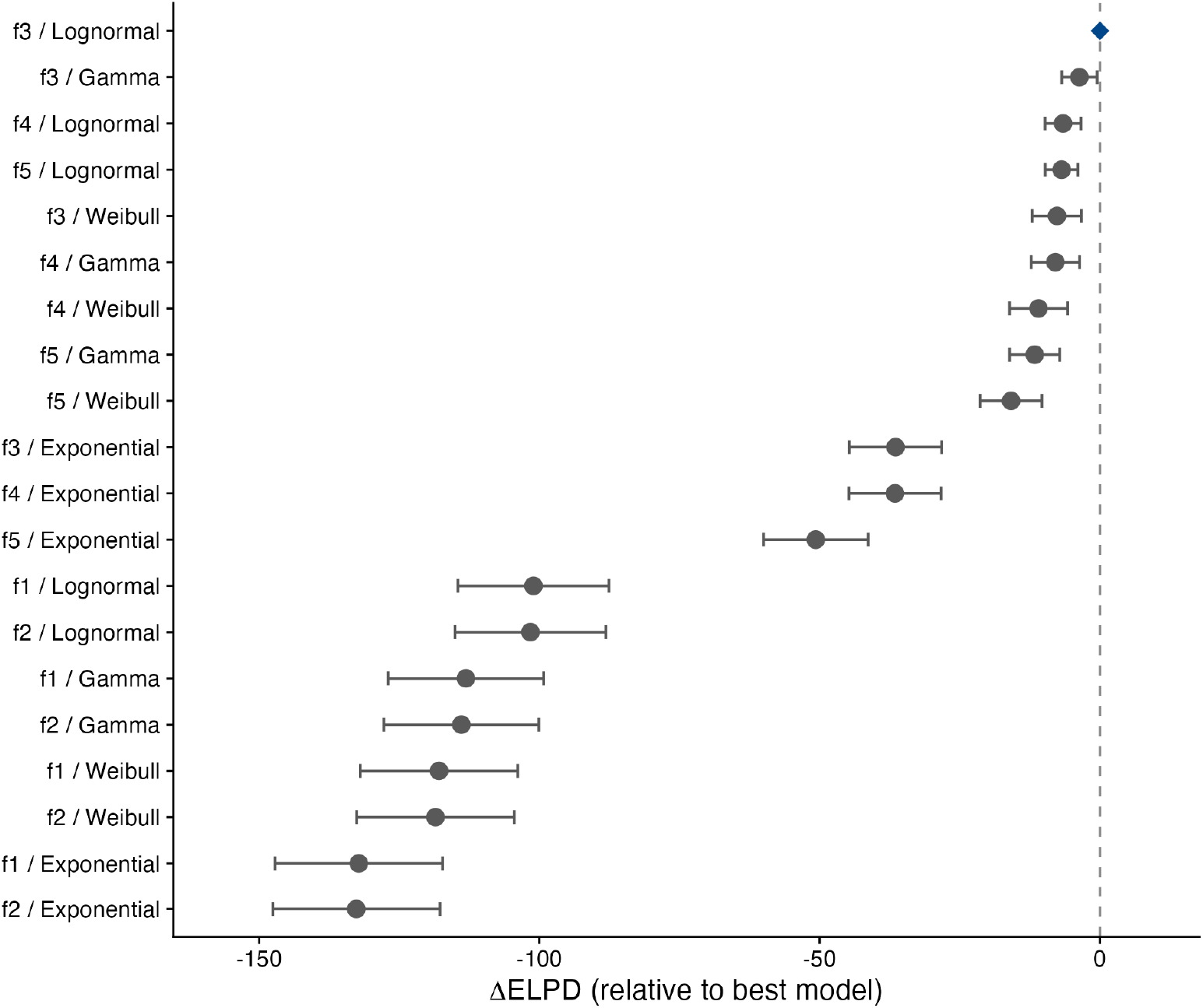
All-model PSIS-LOO comparison ranked by predictive accuracy. ΔELPD relative to the best-performing model (diamond, blue) with *±*1 SE error bars. Models with uncertainty intervals overlapping zero have predictive performance that is not clearly different from the best-performing model. Labels indicate the temperature-response structure and survival distribution.

Within the f3 structure, alternative survival distributions showed comparable predictive performance. The f3 weibull model had overlapping LOO uncertainty intervals with the f3 lognormal model and differed by fewer than 4 ELPD units, indicating limited evidence for improved predictive performance from alternative tail distributions once the temperature-response structure was specified. The lognormal distribution was retained as the final model based on its marginally higher ELPD and consistency with previous Bayesian EIP modelling approaches for arboviruses [6].

The final model was:

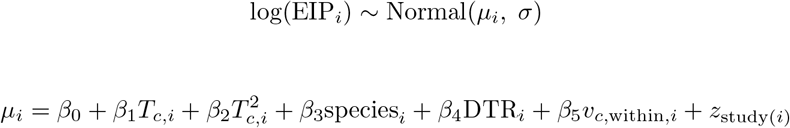

where *i* indexes individual mosquito observations, *T*_*c*_ represents mean-centred temperature, *β*_0_ is the intercept, *β*_1_ *−β*_5_ are regression coefficients, and *z*_study(*i*)_ is the study-level random intercept:

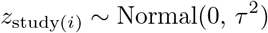

where *τ* represents between-study variation in baseline EIP.

### Temperature Dependence of EIP

Across both vector species, ZIKV EIP declined steeply and non-linearly with increasing temperature, with the greatest reductions occurring between 20°C and 25°C and progressively smaller reductions above 28–30°C (Figure 2, Table 1). In *Ae. aegypti*, median EIP decreased from 59.5 days (95% CrI 27.0–124.0) at 20°C to 15.9 days (7.3–32.2) at 25°C, representing an approximately 73% reduction over five degrees. EIP declined further to 10.2 days (4.6–20.5) at 28°C and 8.4 days (3.8–17.0) at 32°C. Similar estimates at 30°C (8.8 days, 4.0–17.6) and 32°C indicate that additional reductions in ZIKV EIP were limited at higher temperatures within the observed experimental range.

**Fig 2.**
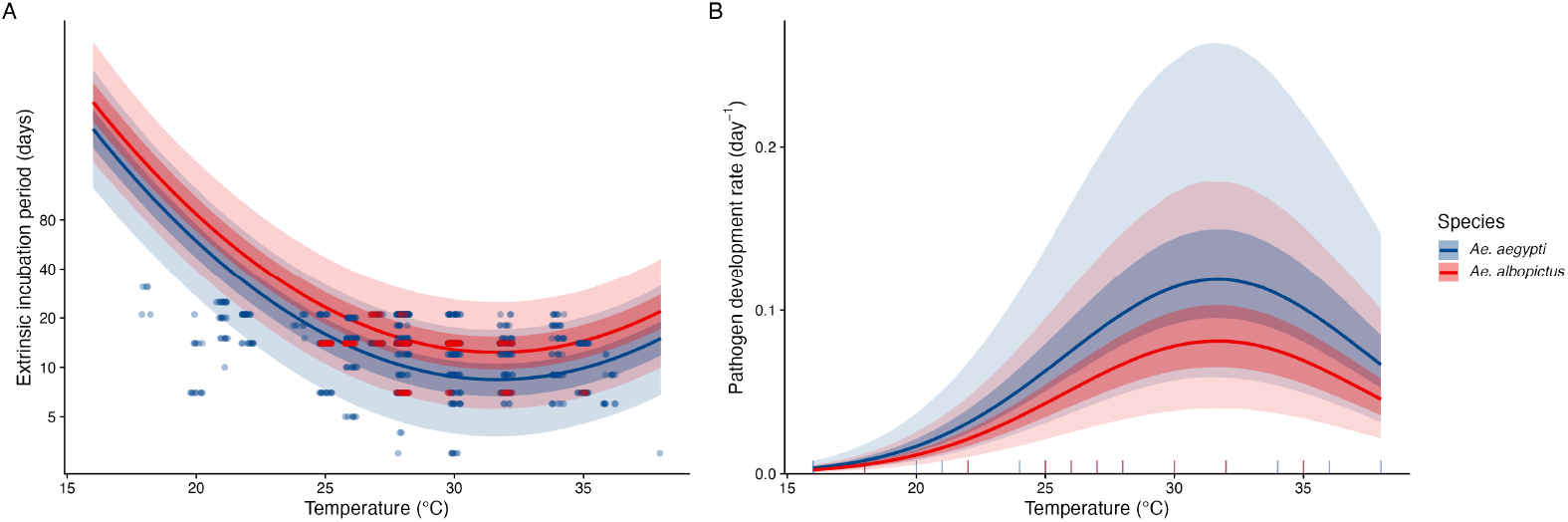
Posterior predicted extrinsic incubation period (EIP; panel A) and pathogen development rate (PDR = 1*/*EIP_50_; panel B) for *Ae. aegypti* and *Ae. albopictus* across experimental temperatures from the final model (f3 lognormal). Lines show posterior median predictions; dark and light ribbons represent 50% and 95% credible intervals. Predictions were generated at DTR = 0 with study-level random effects marginalised. Points in panel A show observed transmission-positive sampling time points (upper bounds of interval-censored observations), jittered horizontally. Rug marks in panel B show the distribution of experimental temperatures contributing to each species.

**Table 1.**
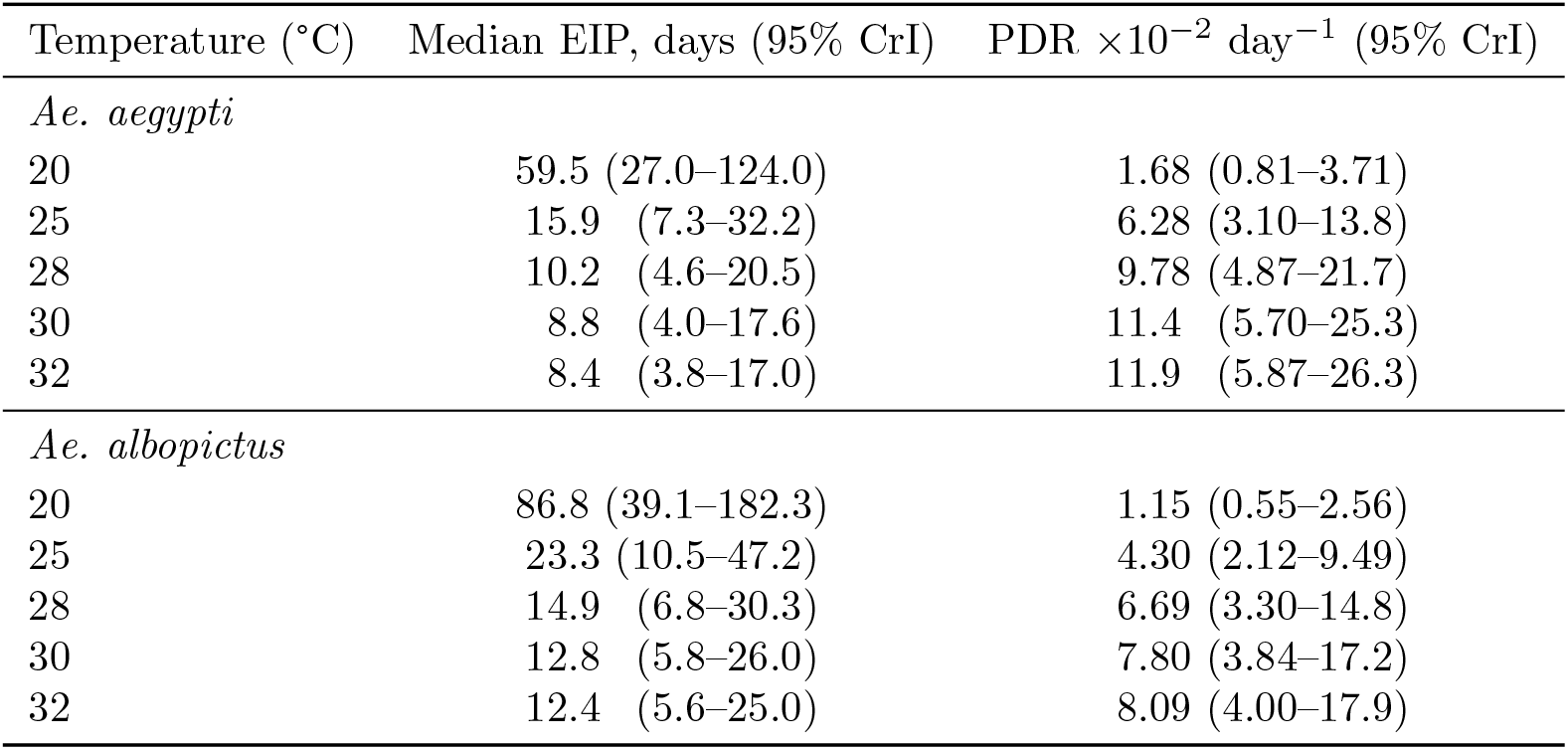
Posterior predicted median EIP and pathogen development rate (PDR) at ecologically relevant temperatures for *Ae. aegypti* and *Ae. albopictus*. PDR = 1*/*EIP_50_. Values in parentheses are 95% credible intervals. Predictions at DTR = 0 with study-level random effects excluded.

**Table 2.** Candidate model structures evaluated for ZIKV extrinsic incubation period. All models included mean-centred incubation temperature 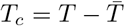 diurnal temperature range (DTR), within-study centred log viral load (*v*_*c,within*_), and a study-level random intercept (1 | ID).

| Model | Temperature structure | Species term | Biological interpretation |
| --- | --- | --- | --- |
| f1 | $T_c$ | + species | Linear temperature response shared between species, with different baseline EIPs |
| f2 | $T_c \times \text{species}$ | interaction | Species-specific linear temperature responses |
| f3 | $T_c + T_c^2$ | + species | Shared nonlinear quadratic temperature response, with different baseline EIPs |
| f4 | $s(T_c, k = 5)$ | + species | Shared flexible nonlinear temperature response, with different baseline EIPs |
| f5 | $0 + \text{species} + s(T_c, \text{by} = \text{species}, k = 5)$ | species-specific smooths | Species-specific flexible nonlinear temperature responses |

*Ae. albopictus* exhibited the same non-linear temperature dependence but consistently longer EIP durations. Median EIP decreased from 86.8 days (39.1–182.3) at 20°C to 23.3 days (10.5–47.2) at 25°C, followed by further reductions to 14.9 days (6.8–30.3) at 28°C and 12.4 days (5.6–25.0) at 32°C (Table 1). Across the full temperature range, *Ae. albopictus* exhibited an EIP approximately 1.5-fold longer than *Ae. aegypti*, consistent with the additive species effect estimated by the final model: the two species differed in baseline EIP while sharing a common temperature response. The reciprocal transformation of EIP to pathogen development rate (PDR = 1*/*EIP_50_) is shown in Figure 2 to facilitate incorporation of these estimates into temperature-dependent vectorial capacity models.

The full posterior-predicted distribution of individual EIP times across temperatures is shown in Figure 3. At lower temperatures, predicted EIP distributions were broad and strongly right-skewed, reflecting greater uncertainty and variability in the timing of acquisition of transmission potential. At warmer temperatures, distributions shifted towards shorter EIP durations and became narrower, indicating more concentrated predictions of transmission timing. This narrowing was more pronounced in *Ae. aegypti* than *Ae. albopictus*, consistent with the shorter predicted EIP duration and lower posterior variability estimated for this species across the temperature range.

**Fig 3.**
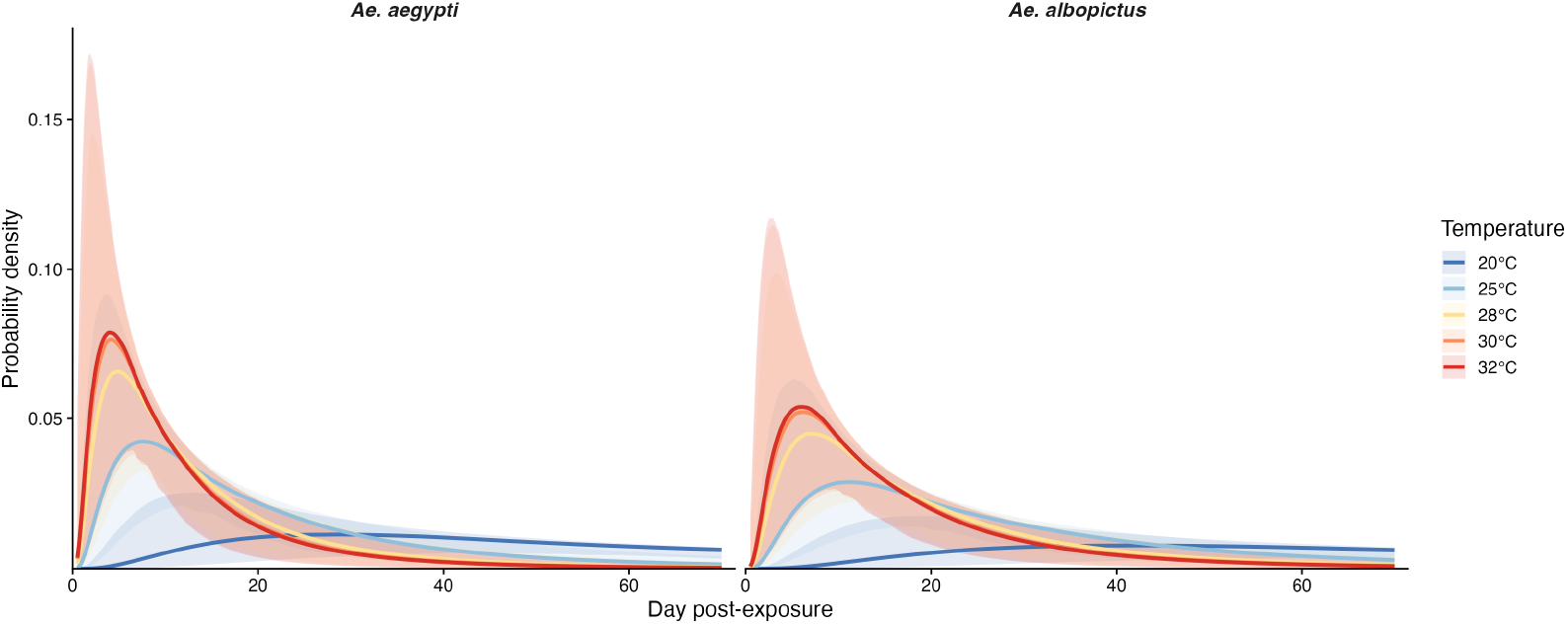
Posterior-predicted probability density of ZIKV EIP at five temperatures for *Ae. aegypti* (left) and *Ae. albopictus* (right). Curves show the lognormal probability density of individual EIP times based on posterior median parameter estimates; shaded ribbons represent 95% credible intervals across posterior draws.

### Sensitivity analyses

The estimated temperature–EIP relationship was robust to both leave-one-study-out and prior sensitivity analyses (Supplementary Index S3, Figures S1 and S2). Removal of individual studies did not materially alter posterior median EIP estimates across the temperature range with substantial experimental coverage (25–32°C). At lower temperatures, where observations were sparse, excluding individual studies resulted in wider credible intervals, reflecting reduced information rather than systematic changes in the estimated temperature relationship. Prior sensitivity analyses showed that posterior predictions were generally stable across alternative prior specifications, with minor differences restricted primarily to the lower temperature range where experimental data were limited.

## Discussion

This study provides the first Bayesian synthesis of ZIKV extrinsic incubation period (EIP) across multiple independent vector competence studies, extending censored survival modelling approaches previously applied to dengue [6], yellow fever [24], and West Nile virus [16] to ZIKV in *Ae. aegypti* and *Ae. albopictus*. Consistent with previous analyses, we find that EIP decreases strongly with increasing temperature and that parametric survival models provide a flexible framework for representing variation in individual transmission timing. The lognormal distribution selected for ZIKV is consistent with the dengue analysis of Chan and Johansson (2012) [6], whereas a Weibull distribution was selected for West Nile virus and yellow fever virus [16, 24]. The identification of right-skewed distributions across these arbovirus systems suggests that parametric survival approaches provide a useful general framework for modelling heterogeneous EIP dynamics.

In contrast to previous studies, we find strong evidence that the ZIKV EIP–temperature relationship is non-linear. Whereas Chan and Johansson (2012) described dengue EIP using a linear temperature relationship on the log scale, and Vollans et al. (2024) found limited support for additional quadratic terms for West Nile virus, the quadratic temperature model substantially improved predictive performance for ZIKV compared with linear alternatives [6, 16]. This indicates that the rate of EIP reduction with increasing temperature changes across the observed thermal range. By pooling data from eight independent studies and two vector species, accounting explicitly for interval censoring at the individual mosquito level, and comparing alternative temperature-response structures, this analysis provides an uncertainty-calibrated characterisation of ZIKV transmission timing across experimentally observed temperatures.

The estimated temperature dependence of ZIKV EIP is consistent with experimental evidence showing that temperature is a major determinant of ZIKV replication and transmission efficiency in *Aedes* mosquitoes [4, 5, 10]. Previous vector competence studies have demonstrated shorter EIPs and increased transmission efficiency across permissive temperature ranges, although estimates vary substantially between mosquito populations, viral strains, and experimental protocols [5, 10, 11]. The predicted EIP values from our model integrate this heterogeneous evidence base and provide a continuous temperature-response function rather than isolated estimates from individual experiments. This distinction is important because small differences in EIP can substantially influence vectorial capacity: mosquitoes must survive the EIP before contributing to onward transmission, meaning that temperature-driven reductions in EIP can disproportionately increase epidemic potential [4]. These temperature-dependent effects are particularly relevant under scenarios of climate change, where shifts in environmental conditions may alter the geographic and seasonal suitability for ZIKV transmission by modifying mosquito suitability, vector competence, and pathogen development rates [27, 28].

Several limitations should be considered when interpreting these estimates. Vector competence experiments rely on destructive sampling, meaning that the exact time at which an individual mosquito becomes infectious is not observed. A mosquito positive for infectious virus at time *t* may therefore have completed its EIP at any point between exposure and sampling. Studies with infrequent sampling schedules provide less precise interval-censored observations, and while the Bayesian framework accounts for this uncertainty, it cannot recover information that was not collected experimentally.

The available evidence for additional covariates was also limited. DTR was included because one study evaluated fluctuating temperatures, whereas most studies used constant-temperature conditions and therefore contributed little variation in this predictor [12]. Consequently, the estimated DTR effect should be interpreted cautiously, and predictions presented here represent constant-temperature conditions (DTR = 0). Similarly, the viral load effect was informed primarily by Chouin-Carneiro et al., the only study testing multiple viral load doses under comparable experimental conditions [13]. Although within-study centring reduced confounding between viral load and study-level variation, the limited number of studies contributing dose-response information restricts generalisation of this effect. Furthermore, differences in reported viral load units across studies (e.g., TCID_50_, PFU, CCID_50_) may introduce additional measurement uncertainty that could not be propagated in the present model.

The selected quadratic temperature relationship represents a compromise between biological realism and model identifiability. Thermal performance curves are often asymmetric, with distinct lower and upper thermal limits; however, the available ZIKV EIP data were insufficient to support more complex temperature-response functions without increasing uncertainty. At the extremes of the observed temperature range ( ≤ 18°C and ≤ 34°C), observations were sparse and predictions should therefore be interpreted cautiously. In addition, the study-level random intercept accounts for unexplained variation in baseline EIP but assumes a shared temperature response across studies and species. Remaining heterogeneity may reflect differences in mosquito population origin, laboratory adaptation, ZIKV lineage, and diagnostic methods, including differences between molecular and culture-based detection assays.

Despite these limitations, this study provides the most comprehensive Bayesian synthesis of ZIKV EIP data currently available. By modelling EIP as an interval-censored time-to-event outcome rather than a single threshold estimate, the approach directly represents uncertainty arising from experimental design and biological variation. The stability of results to removal of individual studies and alternative prior specifications further supports the robustness of the estimated temperature relationship. These posterior estimates provide a mechanistically interpretable and uncertainty-quantified representation of ZIKV EIP that can be directly incorporated into temperature-dependent vectorial capacity and transmission models.

## Conclusion

Understanding how temperature shapes the extrinsic incubation period (EIP) of ZIKV is fundamental for predicting transmission potential and improving projections of how environmental change may influence the geographic and seasonal suitability for ZIKV transmission. By applying a Bayesian interval-censored survival framework to pooled data from eight independent laboratory studies, we demonstrate that ZIKV EIP in both*Ae. aegypti* and *Ae. albopictus* is strongly and non-linearly temperature-dependent, with a quadratic relationship on the log scale providing substantially improved out-of-sample predictive accuracy compared with linear temperature-response models. Our estimates were robust to the removal of individual studies and alternative prior specifications, supporting their use as biologically grounded inputs for temperature-dependent vectorial capacity and transmission models. As climatic conditions change, nonlinear temperature effects on EIP may alter the periods and locations where environmental conditions are suitable for ZIKV transmission. The calibrated, uncertainty-quantified predictions provided here offer a framework for incorporating temperature-dependent viral development into future projections of ZIKV transmission potential under changing climate scenarios.

## Methods

All analyses were conducted in R (R version 4.5.3 (2026-03-11)) [18].

### Data and experimental design

This analysis used data compiled by Hartner et al. (2025) [17]. Authors conducted a systematic review of laboratory studies examining arbovirus infection efficiency, transmission efficiency, EIP, and temperature relationships using studies published between 1932 and May 2025. Transmission potential was assessed by detection of infectious virus in mosquito saliva or salivary glands. The systematic review protocol and data extraction procedures are described in the original publication [17].

Vector competence experiments typically rely on destructive sampling, where cohorts of mosquitoes are sacrificed at predefined time points following exposure to determine whether infectious virus is present in saliva [15]. Consequently, the exact time at which an individual mosquito becomes capable of transmission is not directly observed. Instead, each observation represents an interval- or right-censored survival outcome. For mosquitoes with detectable virus in saliva at time *t*, the EIP is interval-censored between exposure and the sampling time point *t*. For mosquitoes without detectable virus in saliva at time *t*, the EIP is right-censored at *t*, as transmission competency had not occurred by the time of observation. Within a mosquito cohort, the median EIP (EIP_50_) represents the time point at which 50% of infected mosquitoes are predicted to have become infectious.

Here, we focus on ZIKV data from eight *Ae. aegypti* and *Ae. Albopictus* studies [4, 12–14, 19–22]. Only mosquitoes with confirmed infection were used as the at-risk denominator, ensuring that the model estimates the true EIP conditional on infection rather than conflating EIP with vector competence. For studies in which transmission was assessed in only a subset of infected mosquitoes, the number of saliva-positive mosquitoes was estimated by applying the observed transmission proportion among tested mosquitoes to the full infected cohort:

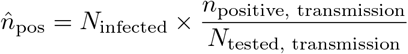

where *N*_infected_ is the total number of infected mosquitoes, *n*_positive, transmission_ is the number of mosquitoes with detectable virus in saliva, and *N*_tested, transmission_ is the number of mosquitoes assessed for transmission. Derived counts were rounded to the nearest integer.

Aggregated study-level observations were expanded into individual mosquito-level records representing either interval-censored transmission events or right-censored non-transmission observations. A small offset of 0.001 days was applied to zero-valued interval lower bounds to ensure compatibility with strictly positive parametric survival distributions. An overview of the final dataset is provided in S1 Appendix.

### Statistical modelling

The interval- and right-censored nature of the experimental observations motivated analysis of ZIKV EIP using a censored time-to-event framework implemented in the Bayesian modelling package brms (version ≤2.21) [23]. Bayesian survival approaches have previously been applied to estimate EIP for vector-borne diseases [6, 16, 24]. Models were compared using approximate leave-one-out cross-validation (LOO-CV) implemented in the **loo** package [25].

Four parametric survival distributions were evaluated: **lognormal, Weibull, gamma**, and **exponential**, consistent with previous comparative analyses of EIP models [6, 16, 24]. Rather than specifying the survival distribution a priori, each distribution was evaluated across five candidate predictor structures (f1–f5; described below), resulting in 20 candidate models.

For observation *i*, the likelihood contribution under parametric distribution *F* ( ·|***θ***) was defined as:

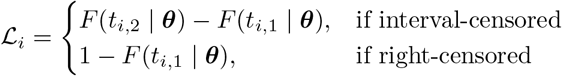

where *t*_*i*,1_ and *t*_*i*,2_ represent the lower and upper bounds of the censoring interval for observation *i*.

### Candidate model structures

Five candidate formula structures were evaluated to represent alternative hypotheses regarding the temperature–EIP relationship and the extent to which this relationship differs between mosquito species. All models included a study-level random intercept (1| ID) to account for unmeasured between-study heterogeneity, including potential differences in study design, virus strains, mosquito rearing conditions, and assay methods. All models also included diurnal temperature range (DTR), mean-centred incubation temperature 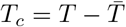 and within-study centred log viral load (*v*_*c,within*_) as covariates. Temperature was mean-centred to improve numerical stability and allow the intercept to represent the expected EIP at the mean observed experimental temperature.

The candidate structures differed in the assumed form of the temperature response and the degree of flexibility allowed between species. Model f1 represents the simplest assumption of a shared linear temperature response with species-specific baseline differences. Model f2 extends this by allowing species-specific linear temperature responses. Model f3 introduces a nonlinear quadratic temperature response while retaining a common temperature relationship across species. Models f4 and f5 use penalised regression splines to relax assumptions about the functional form of the temperature response. Model f4 estimates a shared nonlinear temperature response with species-specific baseline offsets, whereas model f5 allows both baseline EIP and the temperature response function to vary between species.

Penalised regression splines were implemented using the **mgcv** framework with a fixed basis dimension of k=5 [26].

### Viral load and collinearity

Because most studies used a single viral dose within a study, a raw viral load covariate would primarily capture between-study differences in experimental conditions rather than within-study variation in dose response. These between-study differences are not distinguishable from other sources of study-level heterogeneity captured by the random intercept (1| ID). To avoid attributing study-level differences in baseline EIP to viral load, viral load was therefore centred within study:

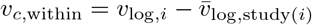

Where 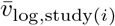 represents the mean log viral load within the study containing observation *i*. This transformation retains only within-study viral dose variation, which was available in Chouin-Carneiro et al. [13], while allowing between-study differences in viral dose and other experimental conditions to be captured by the study-level random intercept. Studies with a single viral load concentration contribute 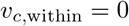

### Prior specification

Priors were informed by published EIP estimates from related arboviruses and plausible ranges for temperature-dependent transmission dynamics. Prior distributions were selected to constrain parameter estimates to biologically realistic values while remaining sufficiently weakly informative given the heterogeneity of the available experimental data. Full prior specifiscations are provided in S2 Appendix.

### Posterior inference and convergence

All 20 models were fitted using four independent Markov chains, each run for 4,000 iterations with 2,000 warm-up iterations, resulting in 8,000 posterior draws per model. The target acceptance probability was set to *δ* = 0.98, with a maximum tree depth of 15.

Convergence was assessed using the potential scale reduction factor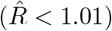, bulk and tail effective sample sizes (*n*_eff_ *>* 400), and divergent transitions. Pareto-*k* diagnostics from PSIS-LOO were inspected to identify influential observations.

### Model selection

All 20 candidate models were compared using Pareto-smoothed importance sampling leave-one-out cross-validation (PSIS-LOO) [25]. The model with the highest estimated expected log predictive density (ELPD) was selected, with the more parsimonious model preferred when competing models showed comparable predictive performance (ELPD difference *<* 2*×* the standard error of the difference).

Predictions were generated over a fine temperature grid spanning the observed experimental temperature range. Population-level predictions excluded study-level random effects. Pathogen development rate (PDR) was calculated as the reciprocal of the posterior median EIP (PDR = 1*/*EIP_50_), with posterior medians and 95% credible intervals (CrI) derived from posterior predictive distributions.

### Sensitivity analyses

Two sensitivity analyses were conducted to evaluate the robustness of conclusions from the final model (f3 lognormal). Full results are provided in S3 Appendix.

#### Leave-one-study-out (LOSO)

To assess the influence of individual studies on the estimated temperature–EIP relationship, the final model was refitted after sequentially removing each study. Posterior predictions were compared across the observed temperature range, with changes in posterior uncertainty assessed relative to the full model. Changes in posterior uncertainty, quantified by the width of the 95% credible intervals across temperatures, were compared between each leave-one-study-out model and the full model.

#### Prior sensitivity

The influence of prior assumptions on posterior predictions was assessed by varying prior scale parameters individually while holding all other priors constant. Prior sensitivity analyses varied prior scale parameters individually, including the intercept, fixed-effect coefficients, and residual variance, while holding other priors constant. For each component, tighter and wider prior variants were evaluated.

## Supporting information

Supplementary Index 1

Supplementary Index 2

Supplementary Index 3

## Supporting information

**S1 Appendix. Characteristics of the studies included in the ZIKV EIP analysis**

**S2 Appendix. Prior Selection** A description of the priors used in the models.

**S3 Appendix. Sensitivity Analysis.**

## Acknowledgments

AMH and CI acknowledge funding from The Federal Ministry of Research, Technology and Space (BMFTR) through the CLIMADEMIC project (funding code 01LN2210A) within the framework of the Strategy Research for Sustainability (FONA). LW declarex no funding. These funders had no role in study design, data collection and analysis, decision to publish, or preparation of the manuscript.

We have read the journal’s policy and the authors of this manuscript have the following competing interests: AMH received additional funding from Gavi, the Vaccine Alliance, the Bill & Melinda Gates Foundation and the Wellcome Trust via the Vaccine Impact Modelling Consortium (VIMC) during the course of the study (grant number INV-034281). These funders had no role in study design, data collection and analysis, decision to publish, or preparation of the manuscript. All other authors declare no competing interests.

## Data availability statement

All data and code used for running experiments, model fitting, and plotting is available on a GitHub repository at XX (will be made available on publishing).

