## Supplementary Index 1 for "The extrinsic incubation period for Zika virus: a Bayesian time delay modelling study"

### S1 Appendix. Characteristics of the studies included in the ZIKV EIP analysis

Table 1 summarises the eight vector competence studies included in the final ZIKV EIP dataset. Together, these studies encompass experiments conducted in *Ae. aegypti* and *Ae. albopictus* across a range of mosquito populations, ZIKV strains, viral inoculation doses, incubation temperatures, sampling schedules, and diurnal temperature regimes. Following data processing described in the main text, aggregated cohort observations were expanded to individual mosquito-level interval- and right-censored observations for Bayesian survival analysis.

**Table 1.** Studies included in the ZIKV extrinsic incubation period analysis.

| Study | Species | Mosquito origin | Strain | Viral load given | Temp. range (°C) | Sampling days | DTR (°C) |
|---|---|---|---|---|---|---|---|
| Chouin-Carneiro 2020 [1] | Ae. aegypti, Ae. albopictus | Brazil | American lineage (BRPE243/2015; GenBank: KX197192) | 10ˆ2–10ˆ6 PFU/mL | 22–28 | 14–21 | 0 |
| Heitmann 2017 [2] | Ae. aegypti, Ae. albopictus | Germany; Italy | FB-GWUH-2016, GenBank: KU870645 | 10ˆ7 PFU/mL | 18–27 | 14–21 | 0 |
| Hernández-Triana 2019 [3] | Ae. aegypti | Cuba | Strain H/PPF/2013; GenBank: KJ77679 | 10ˆ7.2 PFU/mL | 20–25 | 7–21 | 0 |
| Hugo 2019 [4] | Ae. aegypti, Ae. albopictus | Australia | KU365780 | 10ˆ8.8 CCID50/mL | 28 | 3–14 | 0, 7.5 |
| Murrieta 2021 [5] | Ae. aegypti, Ae. albopictus | Mexico; United States | PRVABC59, GenBank: KU501215 | 10ˆ7 PFU/mL | 25–35 | 7–14 | 0, 10 |
| Onyango 2020 [6] | Ae. aegypti, Ae. albopictus | Mexico; United States | HND (2016-19563, GenBank: KX906952) | 10ˆ8.3 PFU/mL | 26–30 | 4–14 | 4 |
| Tesla 2018 [7] | Ae. aegypti | Mexico | MEX1-44 | 10ˆ6 PFU/mL | 16–38 | 3–21 | 0 |
| Winokur 2020 [8] | Ae. aegypti | United States | PRVABC59, PR15, GenBank:KX601168 | 10ˆ5.5 PFU/mL | 18–30 | 3–31 | 0 |

DTR = diurnal temperature range. PFU = plaque-forming units. $CCID_{50}$ = 50% cell culture infectious dose. Temperatures are incubation temperatures following the infectious blood meal.
