## Supplementary Index 2 for "The extrinsic incubation period for Zika virus: a Bayesian time delay modelling study"

### S2 Appendix. Prior specification

Weakly informative priors were specified for all model parameters to regularise estimation while remaining sufficiently broad for inference to be driven primarily by the data. Prior locations were informed by previous Bayesian analyses of arboviral extrinsic incubation periods [1–3] together with biologically plausible ranges for ZIKV EIP. Model-specific priors are summarised in Table 1.

For models including quadratic temperature effects, a tighter prior was assigned to the quadratic coefficient to discourage implausibly large curvature outside the observed temperature range. Species-specific spline models (f5) estimated separate baseline intercepts for each mosquito species, whereas all other models estimated a single global intercept. Distribution-specific parameters were assigned weakly informative priors appropriate for each survival family. The influence of prior specification on posterior inference was assessed through the prior sensitivity analysis described in the main text.

**Table 1.** Summary of prior distributions used across candidate models.

| Parameter | Prior | Purpose |
|---|---|---|
| Intercept | Normal(2.3, 0.77) | Baseline log-EIP |
| Linear temperature | Normal(−0.07, 0.15) | Temperature effect |
| Quadratic temperature | Normal(0, 0.005) | Nonlinear curvature |
| Species, DTR, viral load | Normal(0, 0.23) | Fixed effects |
| Spline SD | Normal(0, 0.5) | Smoothness penalty |
| Lognormal $\sigma$ | Normal(log(0.7), 0.5) | Residual variation |
| Gamma shape | Normal(log(1.5), 0.5) | Shape parameter |
| Weibull shape | Normal(log(1.5), 0.5) | Shape parameter |
