## Supplementary Index 3 for "The extrinsic incubation period for Zika virus: a Bayesian time delay modelling study"

### S3 Appendix. Sensitivity Analysis

The final model (f3_lognormal) was refitted after sequentially removing each study. Posterior predictions from each leave-one-study-out model are shown in Figure 1. Increased uncertainty at lower temperatures reflects the limited number of observations contributing information in this region of the temperature range. Predictions at warmer temperatures (25–32°C), where data density was greater, showed substantially more consistent estimates across leave-one-study-out fits.

Prior sensitivity analyses were conducted for the final model (f3_lognormal) by varying the intercept, fixed-effect, residual variance, and between-study variance priors individually while holding all other priors constant. Posterior temperature–EIP relationships under alternative prior specifications are shown in Figure 2.

**Fig 1. Leave-one-study-out (LOSO) sensitivity analysis of the final model (f3_lognormal).** Posterior predicted median ZIKV extrinsic incubation period (EIP) across temperature after sequential exclusion of individual studies. The black line and grey ribbon show predictions from the full model, while coloured lines show posterior median predictions from models refitted after omitting one study at a time. Predictions are shown for *Ae. aegypti* with study-level random effects marginalised, DTR = 0, and within-study centred viral load set to zero.s

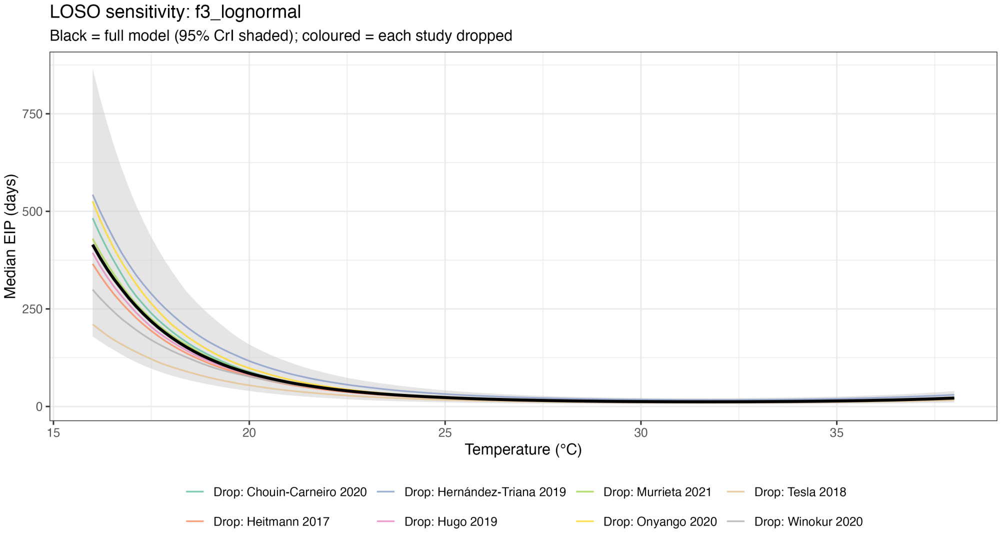

**Fig 2. Prior sensitivity analysis for the final model (f3_lognormal).** Posterior predicted median ZIKV extrinsic incubation period (EIP) across temperature under alternative prior specifications. In each panel, one prior component was varied while all others were held at their baseline values. Black solid lines denote the baseline prior, blue dashed lines tighter priors, and red dotted lines wider priors; shaded ribbons indicate 95% credible intervals. Predictions are shown for *Ae. aegypti* with study-level random effects marginalised, DTR = 0, and within-study centred viral load set to zero.

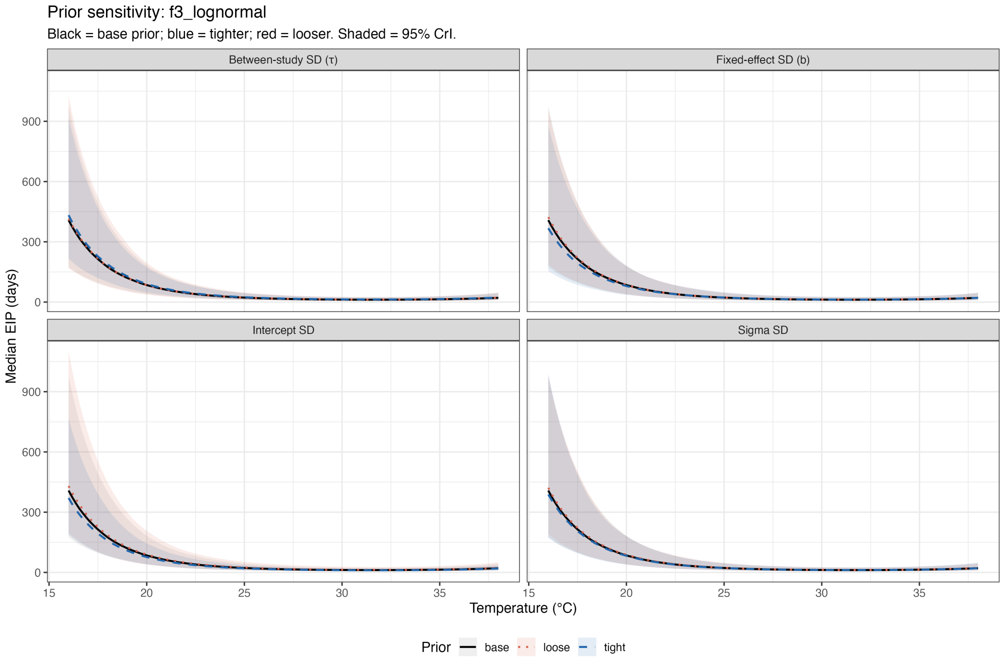
